# Intermanual transfer of visuomotor learning is facilitated by a cognitive strategy

**DOI:** 10.1101/2021.10.12.464030

**Authors:** Jack De Havas, Patrick Haggard, Hiroaki Gomi, Sven Bestmann, Yuji Ikegaya, Nobuhiro Hagura

**Affiliations:** NTT Communication Science Laboratories, Japan; Institute of Cognitive Neuroscience, University College London, UK; Center for Information and Neural Networks, National Institute for Information and Communications Technology, Osaka, Japan; UCL Queen Square Institute of Neurology Department of Clinical and Movement Neurosciences, University College London, UK; Wellcome Centre for Human Neuroimaging, University College London, UK; Graduate School of Pharmaceutical Sciences, Faculty of Pharmaceutical Sciences, The University of Tokyo, Tokyo, Japan; Graduate School of Frontier Biosciences, Osaka University, Osaka, Japan

**Keywords:** visuomotor adaptation, cognitive strategy, intermanual transfer, visuomotor gain, visual feedback

## Abstract

Humans continuously adapt their movement to a novel environment by recalibrating their sensorimotor system. Recent evidence, however, shows that explicit planning to compensate for external changes, i.e. a cognitive strategy, can also aid performance. If such a strategy is indeed planned in external space, it should improve performance in an effector independent manner. We tested this hypothesis by examining whether promoting a cognitive strategy during a visual-force adaptation task performed in one hand can facilitate learning for the opposite hand. Participants rapidly adjusted the height of visual bar on screen to a target level by isometrically exerting force on a handle using their right hand. Visuomotor gain increased during the task and participants learned the increased gain. Visual feedback was continuously provided for one group, while for another group only the endpoint of the force trajectory was presented. The latter has been reported to promote cognitive strategy use. We found that endpoint feedback produced stronger intermanual transfer of learning and slower response times than continuous feedback. In a separate experiment, we confirmed that the aftereffect is indeed reduced when only endpoint feedback is provided, a finding that has been consistently observed when cognitive strategies are used. The results suggest that intermanual transfer can be facilitated by a cognitive strategy. This indicates that the behavioral observation of intermanual transfer can be achieved either by forming an effector-independent motor representation, or by sharing an effector-independent cognitive strategy between the hands.

**New and noteworthy:** The causes and consequences of cognitive strategy use for motor learning are poorly understood. We tested whether a visuomotor task learned using a strategy generalizes across effectors. Visual feedback was manipulated to enhance the use of a cognitive strategy. Learning using a cognitive strategy for one hand transferred to the task performed by the un-learned hand. Our result suggests that intermanual transfer can also result from a common cognitive strategy used to control both hands.

## Introduction

When hitting a tennis ball on a windy day, you might aim slightly to the side of where you want the ball to land in order to take the direction of the wind into account. As such, humans can explicitly shift the aim of their actions to compensate for external perturbations; known as a cognitive strategy. Although error-based motor learning has traditionally been considered a single implicit sensorimotor recalibration process (Kawato 1999), recent work has described the contribution of such cognitive strategies to motor learning (Krakauer et al. 2019; Miyamoto et al. 2020).

Cognitive strategies differ from motor adaptation in terms of how and where in the brain they are implemented (Jahani et al. 2020; Serrien et al. 2002). They also likely differ in terms of how sensory feedback is processed, with cognitive strategies producing learning that weights performance error above sensory prediction error to a greater extent than learning by adaptation (Taylor and Ivry 2012). Learning using cognitive strategies and motor adaptation overlap throughout sensorimotor tasks (McDougle et al. 2015), but can be separated by manipulating task instructions (Mazzoni and Krakauer 2006; Taylor and Ivry 2011). In this study, we investigate how cognitive strategy use generalize across effectors in motor learning, by examining the intermanual transfer of motor learning.

How motor adaptation tasks learned on one hand transfer to the other has been extensively studied (Anguera et al. 2007; Imamizu and Shimojo 1995; Wang and Sainburg 2003, 2006). This intermanual transfer has been traditionally ascribed to motor adaptation happening in each hemisphere (Ruddy and Carson 2013), however, whether a cognitive strategy can facilitate intermanual transfer is still under debate. Some studies have reported that the use of a cognitive strategy during motor adaptation tasks can facilitate intermanual transfer (Malfait and Ostry 2004; Werner et al. 2019), whereas others have not (Taylor, Wojaczynski, and Ivry 2011; Wang, Joshi, and Lei 2011; Wang, Lei, and Binder 2015). In these studies, cognitive strategy use has been promoted by introducing an abrupt change in the perturbation (i.e. sudden introduction of the visuomotor rotation or force field), purposefully making the participants aware of the perturbation. However, this method may potentially induce inter-individual variability in cognitive strategy use, depending on the size of the change and differences in individual sensitivity to that change (Werner, Strüder, and Donchin 2019).

In this study, we promote the use of cognitive strategy during motor learning by showing only the endpoint of the action (Endpoint Feedback; EPF), as opposed to showing the feedback continuously throughout the action (Continuous visual feedback; CVF). CVF provides both visual sensory prediction errors and visual performance errors relating to the entire action. EPF conversely, involves a single visual performance error signal pertaining to goal completion. Since cognitive strategies may preferentially weight performance error, we predict that restricting visual feedback to a salient performance error signal may shift the means of task learning away from motor adaption and towards strategy use. Indeed, aftereffects upon the removal of a visuomotor perturbation, a hallmark of motor adaptation, are attenuated by EPF relative to CVF (Hinder et al. 2008; Barkley et al. 2014; Taylor et al. 2014).

In the task, participants isometrically and ballistically exerted force on a gripped handle to control a visual bar-height on screen to reach a target height. After a baseline phase the visuomotor gain (force to bar-height transformation) increased, requiring participants to modify their motor command in order to maintain performance. A 2×2 across-subjects factorial design was used for the first experiment, with factors of visual feedback (EPF vs. CVF) and perturbation schedule (Abrupt vs. Gradual increase of visuomotor gain), conceptually mimicking previous studies (Werner, Strüder, and Donchin 2019, Taylor, Wojaczynski, and Ivry 2011; Wang, Joshi, and Lei 2011; Wang, Lei, and Binder 2015).

We first assessed whether EPF and abrupt gain change promoted cognitive strategies by examining reaction times. Verbal instructions to use cognitive strategies, tasks showing only EPF, and tasks where perturbations are changed abruptly, all exhibit slow response times (Benson et al. 2011; Saijo and Gomi 2010). Additionally, limiting response times reduces strategic learning, as evidenced by increased aftereffects (Haith et al. 2015). Thus, if EPF and Abrupt gain change do promote strategy use they should be associated with prolonged RT. Second, we examined the transfer rate of a gain change learned with the right-hand to the left hand. Since planning based on performance error is computed in target space (Schween et al. 2020), e.g. to aim more to the right than the target is located, such strategies should be applicable for controlling either hand. Thus, cognitive strategy use should facilitate intermanual transfer. Finally, in a separate experiment, we provided independent evidence that the type of visual feedback provided in our current force production task can indeed promote strategy use, by showing that this factor influences the size of aftereffects, consistent with previous reports (Benson et al. 2011; Haith et al. 2015; Morehead et al. 2015).

## Materials and Methods

### Equipment

Participants were seated and held a plastic handle (aligned to midline, navel height) in a power grip. The handle was instrumented with force sensors, which consisted of an optical strain gauge composed of a digital fiber sensor (FS-N10; Keyence corp.) and a limited-reflective fibre unit (FU-38; Keyence corp.) (Fujiwara et al. 2017). Participants arms were pronated and attached to custom built forearm restraints, which consisted of moulded plastic with Velcro straps at either end (see Fig. 1.). The restraints slotted into adjustable runners attached to a solid wooden board, which allowed rapid arm switching during the task. The force data from the handle was processed online by the connected PC for online presentation of the force (sample rate = 100 Hz). Experimental stimuli were created using Matlab (2017) with Psychophysics Toolbox extensions (Brainard 1997; Pelli 1997) and were presented via a flat screen monitor (27 inch LCD, 1440 × 900 pixels resolution pixels, 60 Hz refresh rate) positioned 40 cm in front of participants.

**Figure 1.**
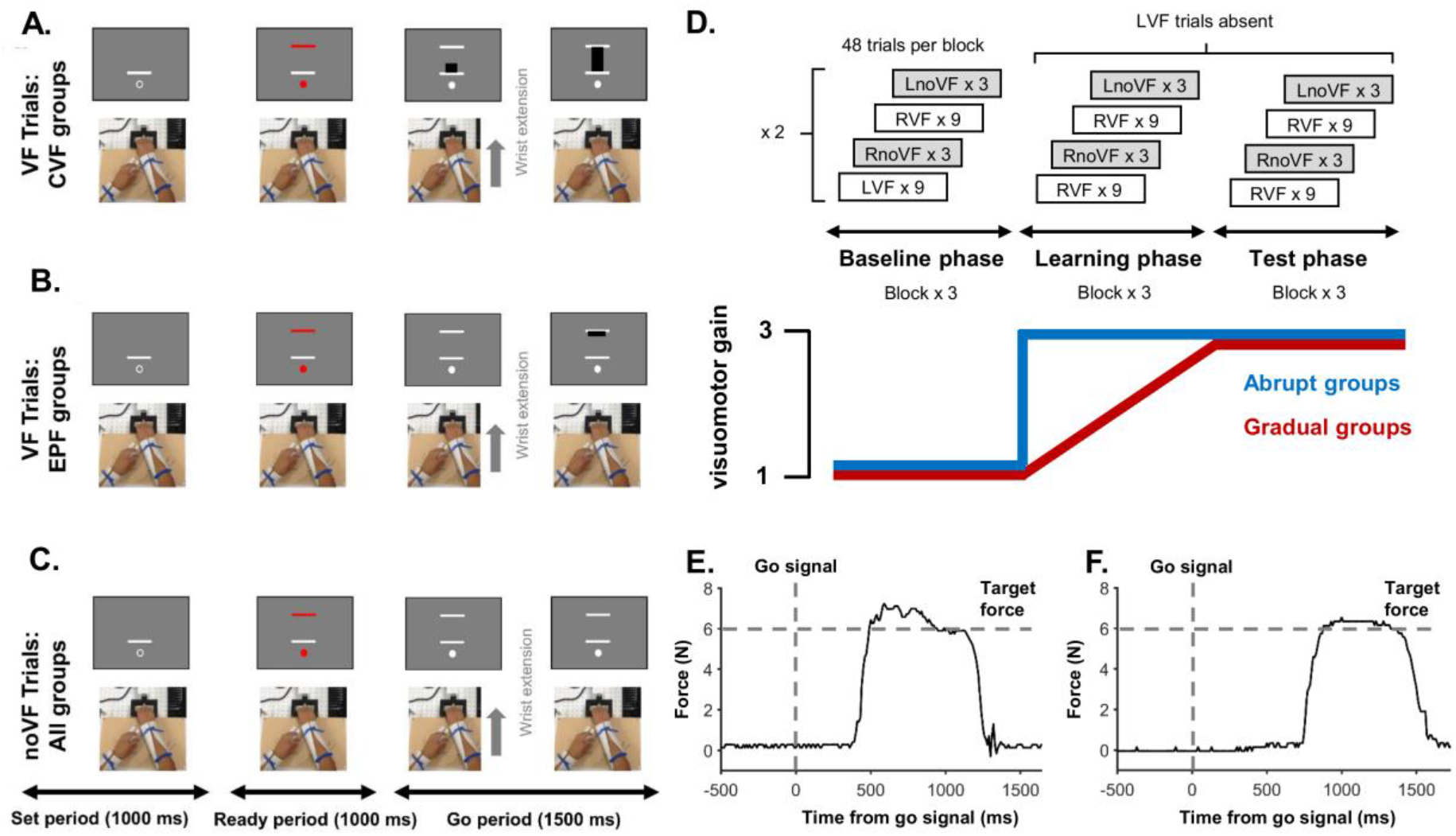
Task structure and single trial results. **A.** Visual feedback trials in CVF groups, where isometric wrist force was continuously shown on screen as the height of a black bar. **B.** For EPF groups, during visual feedback trials force was displayed as a static black bar, which appeared once the wrist extension was completed. **C.** ‘No visual feedback trials’ were identical for all groups and involved participants making wrist extensions of appropriate strength in the absence of any visual feedback. **D.** The experiment had a Baseline, Learning and Test phase, each with 3 blocks of 48 trials. In the Baseline phase participants alternated between sets of 9 ‘visual feedback trials’ and 3 ‘no visual feedback trials’, using either the right or left hand (pseudorandomised). During the Learning phase, visuomotor gain increased from 1 to 3, either abruptly (abrupt gain change groups), or via linear increments across visual feedback trials (gradual gain change groups). Left hand visual feedback trials were absent during the Learning and Test phase, meaning that the gain change was only experienced directly when using the right hand. **E.** A single representative right hand trial from a participant in one of the CVF groups, showing force increase towards the visual target in response to the go signal. **F.** A representative right hand trial from an EPF group participant, showing force increase towards the visual target in response to the go signal.

### Participants

A total of 58 people participated in Experiment 1, of which 2 were excluded for failing to comply with the task, leaving 14 participants per group. CVF Abr: n = 14, Females = 7 (age Mn = 23.8, SD = 4.8). CVF Grd: n = 14, Females = 6, (age Mn = 25, SD = 5.6). EPF Abr: n = 14, Females = 5 (age Mn = 23.4 SD = 3). EPF Grd: n = 14, Females = 5 (age Mn = 24.4 SD = 6.3).

A total of 33 people participated in the Experiment 2. Four participants were excluded from the analysis, two of which were due to mechanical issues and two of which were due to a failure to comply with task instructions. This left 14 participants in the CVF group (Females = 8, age Mn = 22.6, SD = 1.5) and 15 participants in the EPF group (Females = 7, age Mn = 21.3, SD = 1.8), none of whom had participated in Experiment 1.

Both experiments were undertaken with the understanding and written consent of each participant in accordance with the Code of Ethics of the World Medical Association (Declaration of Helsinki), and with approval of the NICT ethical committee. No adverse events occurred during either experiment.

### Procedure

#### Experiment 1

The task was to control the level of force exerted on a handle to reach a target level. The height of the bar on the monitor served as the level of exerted force, and in each trial, participants were asked to set the height of the bar to the target level by isometrically and ballistically exerting force on the handle. The task started with the baseline phase, followed by a learning phase and then the test-phase. After the baseline phase, participants had to adapt to a 3x increase in visuomotor gain in the learning phase (i.e. the same amount of force applied to the handle during the baseline would produce 3x as much bar-height on screen). The increased gain remained stable during the test phase.

Each trial began with participants viewing a white open circle positioned under a white line while holding the handle in their relaxed position (Fig. 1.A-C.). The circle served as a fixation point, and the white bar indicated the baseline force level; the force level when the participants did not intentionally exert force on the handle. After 1000ms, the fixation circle was filled with red and a red target force line appeared at one of three equi-spaced locations above the baseline. Each line height corresponded to three different force levels (3, 6, 9N during baseline phase, 1, 2, 3N during test phase). Participants prepared their response (1000ms) until the central fixation circle and the target force line turned from red to white (Go signal). In response to the Go signal, participants executed isometric wrist extensor contractions of appropriate strength as quickly as possible.

The experiment was designed as a 2 (visual feedback type) x 2 (perturbation schedule) between-subject factorial design, where 4 different combination of factors were assigned to 4 different group of participants.

For the factor of visual feedback type, in one condition, the amount of vertical force exerted on the handle was continuously presented on screen as the height of a solid black bar (Continuous Visual Feedback; CVF) (Fig. 1. A.). In the other condition, feedback was provided instead via a solid black line indicating the force level at the point in time when force velocity had reached its peak (End Point Feedback; EPF) (Fig. 1. B.). For the factor of perturbation schedule, in one condition, the visuomotor gain increased by 3x abruptly at the first trial of the learning phase (Abr). In the other condition, the gain increased gradually (linearly) over the course of learning phase (Grd).

The experiment began with 4 practice blocks (2 blocks per hand) of 48 trials with visual feedback (VF). The baseline phase (Fig 1. D.) consisted of 3 blocks of 48 trials. Each block consisted of 4 iterations of 9 VF trials (3 x low, medium and high force targets, randomised) followed by 3 trials without visual feedback of the exerted force level (noVF) (1 x low, medium and high force targets, randomised). Two of the four sets of 9 VF trials used the right hand (RVF) and two used the left hand (LVF). Likewise, two of the four sets of 3 noVF trials used the right hand (RnoVF) and two used the left hand (LnoVF). In total there were 54 RVF trials, 54 LVF trials, 36 RnoVF trials and 36 LnoVF trials in the baseline phase. Hand order within each block was randomised.

Learning and test phase each had 3 blocks of 48 trials, consisting of similar types of trials as the baseline phase. However, the LVF trials were replaced with the RVF trials, thus, both phases had a total of 108 RVF trials, 36 RnoVF trials and 36 LnoVF trials. This was to prevent any visual error-based learning from occurring for left hand trials while the right hand adapted to the change in the visuomotor gain. Therefore, any visual gain learning observed in the LnoVF trials could be attributed to learning transferred from the right hand. Participants had a 2-minute rest after every task block. The experiment lasted 1.5hrs.

#### Experiment 2

The goal of experiment 2 was to establish whether EPF produced smaller aftereffects than CVF in our force production task, since previous literature using reaching movements suggested that strategy use causes reduced aftereffects. The force generation task in Experiment 2 was the same as Experiment 1. Once again participants exerted force on the handle to reach the same visual targets on screen. But here participants only used their right hand to respond throughout the experiment, and gain changes were identical across the two visual feedback groups (Fig. 3. A.) After a practice block, participants performed 2 blocks (45 trials per block) with a visuomotor gain of 3 (baseline phase) followed by 2 blocks in which the gain gradually decreased to 1. Then followed 4 blocks in which the gain was fixed at 1. The final two blocks with gain fixed to 1 was defined as the test phase, which was used to assess how well participants had learned the gain change. All trials up to this point only included visual feedback trials.

After these blocks of learning the decreased gain from the baseline, on the 10^th^ trial of block 9, the gain suddenly changed back to 3. After this sudden gain change a noVF trial was presented on every third trial for the remainder of block 9 and the entirety of block 10 (total = 27 noVF trials), with the other trials being VF trials (total = 54 trials). These noVF trials, consistent with previous studies (Bond and Taylor 2015; Taylor et al. 2014), were used to assess the size of the aftereffect in the two visual feedback groups (CVF vs EPF). The experiment lasted 1 hour.

### Analysis

#### Experiment 1

For every trial, the time series of the force profile was low-pass filtered using Butterworth filter (5Hz) and the force velocity was calculated. In the CVF groups, response force for each trial was the point at which the force stopped increasing and stabilised, which was determined by taking the point at which the force velocity fell below 10% of the peak force velocity for that trial. This corresponded to what participants attended to on screen and were told would be used to judge their performance accuracy. In the EPF groups, the response force for each trial was the force at the point in time when the force velocity reached its peak, since this corresponded to the feedback presented on screen. In all groups response force was multiplied by the visuomotor gain to transform the force to the visual metrics (i.e. bar height in different gain conditions). This value was transformed into an absolute difference from the target value (absolute error ratio) using the following equation;

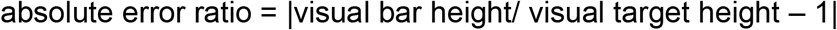

Here, an absolute error ratio of 0 indicates that the force produced was identical to the target level. For no visual feedback trials (noVF), to correct for force drifts before movement onset the data was baseline corrected by subtracting the mean force level during the ready period from the final force level, prior to calculating the absolute error ratio.

Transfer percentages, used to assess intermanual transfer of learning, were calculated from the absolute error ratios in the LnoVF and RnoVF conditions in the following manner for each participant. First the intermanual error ratio was calculated using the absolute error ratio on LnoVF and RnoVF trials;

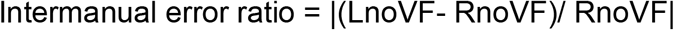

Thus, if the absolute error ratio was equivalent for both hands, the *intermanual error ratio* would be 0, while if it was 3 x larger on the left hand the *intermanual error ratio* would be 2. To make this value more intuitive, the *transfer percentage* was then generated by calculating the *intermanual error ratio* as a percentage of the maximum intermanual error ratio during the test phase, which was 2 (i.e. an intermanual error ratio of 2 is equivalent to an error 3 x larger on the left than the right hand, because none of the 3 x gain increase had been transferred).

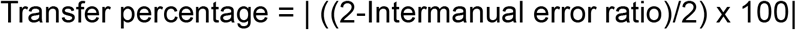

A transfer percentage of 100% therefore meant that the LnoVF absolute error ratio was the same as the RnoVF absolute error ratio, while a transfer percentage of 0% meant the LnoVF absolute error ratio was 3 x larger than the RnoVF absolute error ratio. It should be noted that by setting an upper limit on the error, we are simply normalising to this upper limit. Participants were free to exceed this limit, meaning that *Transfer percentages* can be below 0% or above 100%. The specific value used to define the maximum possible error does not change the results of the statistical tests and is used for display purposes. We used the value of 2 because this was the maximum expected error in the test phase (i.e. if no transfer occurred), and because it approximated the largest errors participants made during the practice session, before the baseline gain settings were learned.

Learning percentages for RVF and RnoVF trials were calculated separately in the same manner, in each case using the mean absolute error ratio at baseline and test.

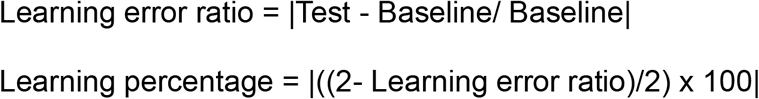

Group differences in transfer percentage at baseline and test, RVF learning percentage at test, and RnoVF learning percentage at test were all assessed using 2 x 2 between subject’s ANOVA, with factors of visual feedback type (CVF vs EPF) and perturbation schedule (Gradual vs Abrupt). We also calculated transfer and learning percentages throughout the experiment by applying the above formulae to every trial. These values were smoothed for display purposes via averaging within a 5-trial moving window.

Reaction times (RT) were calculated for every trial by taking the point in time after the go signal where the force level rose above 4x the SD of the force during the ready period. Mean RT at baseline and test were compared across groups using a 2 x 2 x 2 mixed ANOVA, with the within subject’s factor of phase (baseline vs test) and the between subject’s factors of visual feedback type and perturbation schedule. Trial level RT data was also smoothed for display purposes via averaging within a 5-trial moving window.

Trials were automatically rejected from the analyses based on absolute error ratio if during the 1500ms go period, the participant failed to increase force above 10% of the target force level for that trial. We also rejected trials where peak force velocity occurred after the go period (i.e. late responses > 1500ms). Trials were rejected from the RT analyses if force increases were detected after the go period (late responses > 1500ms) or if RT was classified as being <100ms. There were no significant differences between the CVF and EPF groups in terms of the mean percentage of trials rejected per participant from the error ratio analyses (Mn = 8.77%, SD = 4.96% vs Mn = 11.58%, SD = 7.34%; t(54) = −1.649, p = 0.105) or the RT analyses (Mn = 15.28%, SD = 9.09% vs Mn = 19.96%, SD = 12.44%; t(54) = −1.58, p = 0.12).

#### Experiment 2

RT and absolute error ratio were calculated in the same manner as Experiment 1, as were the learning percentages for RVF trials. We used the same trial exclusion criteria as Experiment 1. There were no significant differences between the CVF and EPF groups in terms of the mean percentage of trials rejected in each participant from the error ratio analyses (Mn = 1.02%, SD = 0.64% vs Mn = 0.81%, SD = 0.9%; t(27) = 0.685, p = 0.499) or the RT analyses (Mn = 10.94%, SD = 14.02% vs Mn = 4.79%, SD = 6.99%; t(27 = 1.511, p = 0.142).

To specifically to assess the size of the aftereffect, we analysed the signed error ratio (i.e. error ratio calculated without converting to absolute values). This was done because during the aftereffect phase the gain suddenly increased from 1 to 3 and we were interested in the degree to which participants overshot the target force level, since this would reflect the degree to which they had adapted to the lower gain setting during the learning and test phases. We determined the size of the aftereffect for each participant for both VF and noVF trials by subtracting the mean signed error ratio during the baseline phase.

We compared mean RT throughout the entire experiment across the two visual feedback groups. We also specifically assessed whether the gradual gain change interacted with group by comparing RT change (test – baseline) in each visual feedback group, and whether the sudden gain change before the aftereffect phase interacted with group by comparing RT change (aftereffect RVF RT – test RVF RT) in each visual feedback group. Independent samples t-tests were conducted on all the variables of interest. Data were smoothed for presentation purposes in the same manner as Experiment. 1.

## Results

### Reaction times were slower for EPF trials

Cognitive strategy use has been associated with prolonged reaction times, possibly due to an increased planning load (Haith et al. 2015; Saijo and Gomi 2010). We found that RT for RVF trials was longer for the EPF groups than the CVF groups throughout the experiment (Fig. 2. D.). This manifested as a significant main effect of visual feedback type when baseline and test phases analyzed for all 4 groups (F(1,52) = 7.041, p = 0.011). It also held when the baseline (F(1,52) = 5.761, p = 0.020), η^2^ = 0.1) and test (F(1,52) = 7.061, p = 0.010, η^2^ = 0.12) phases were analysed separately. There was a trend towards RT getting faster from baseline to test (Main effect of phase: F(1,52) = 3.587, p = 0.064), but no significant interaction between visual feedback type and phase (F(1,52) = 0.032, p = 0.858). Therefore, participants responded more slowly when EPF was available on visual feedback trials throughout the task, possibly by incorporating a cognitive strategy for movement planning.

**Figure 2.**
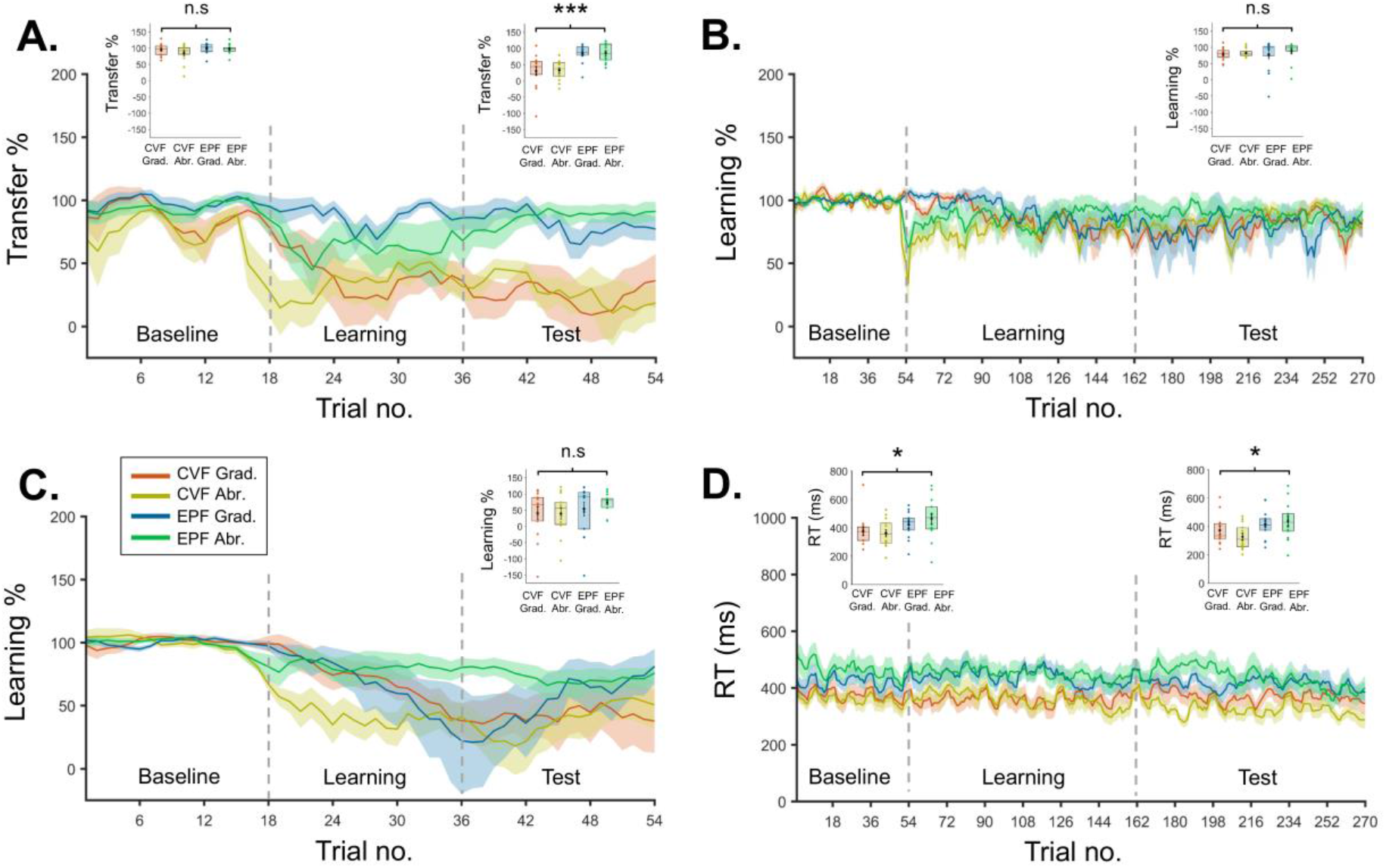
Experiment 1. Transfer, Learning and RT results. **A.** Percentage transfer between RnoVF and LnoVF in each group across the entire experiment. In the EPF groups the amount of transfer returned to around 85% during the learning phase, but in the CVF groups remained around 30% through to the end of the test phase. Insert box plots show that mean transfer % at baseline did not differ across groups, but was significantly higher in the EPF groups than the CVF groups during the test phase (n.s.= not significant, *** p <0.001). **B.** Percentage learning of gain change relative to baseline performance on RVF trials in each group across the entire experiment. Insert box plot shows that learning rates did not significantly differ across groups during the test phase (n.s. = not significant). **C.** Percentage learning of gain change relative to baseline performance on RnoVF trials in each group across the entire experiment. Insert box plot shows that the mean performance at test did not differ across groups (n.s. = not significant). **D.** Response times on RVF trials for all groups across the entire experiment. RT was slower in EPF compared to CVF groups at baseline and test (* p <0.05).

### Greater intermanual transfer of learning for end point feedback

Our main interest was whether a putative shift towards a cognitive strategy has any influence on the intermanual transfer of visuomotor adaptation. We calculated the transfer percentage, which was the ratio between the absolute error ratio of left and right no visual feedback trials, expressed as a percentage of the maximum error. In the learning and test phase, only the right hand was exposed to the perturbation (i.e. visual feedback), but not the left. A transfer percentage close of 100% indicates comparable performance on both hands in the absence of visual feedback, i.e. that all learning on the right hand has been transferred to the left. A transfer percentage of 0% means no learning has been transferred (LnoVF error is 3x larger than RnoVF), while 50% indicates half the learning has been transferred (LnoVF error is 2x larger than RnoVF).

Transfer percentages at test in the EPF groups were higher than those in the CVF groups (EPF Grad. Mn = 84.64%, EPF Abr. Mn = 85.56% vs. CVF Grad. Mn = 30.5%, CVF Abr. Mn = 32.43%; Fig. 2. A. right box plot). ANOVA performed between groups revealed that there was a significant main effect of visual feedback group on the transfer percentage at test (F(1,52) = 31.194, p <0.001, η2 = 0.37). As can be seen from the trial level analysis (Fig. 2. A.), transfer percentages in the EPF groups during the learning phase showed some improvement until reaching a plateau around the start of the test phase. Conversely, transfer percentages in CVF groups were lower, and plateaued earlier during the learning phase.

The transfer results were not due to baseline differences in left and right hand performance when the visual feedback was absent. Baseline transfer percentages were close to 100% in all groups and did not significantly differ from one another (F(1,52 = 2.95, p = 0.092; Fig. 2. A. left box plot).

### Visual gain change was successfully learned for all groups

To determine that the gain change was successfully learned by all participants we calculated the learning percentage on RVF trials, which was the ratio between the absolute error ratio at a given point in time and the absolute error ratio at baseline, expressed as a percentage of the maximum error during the test phase (i.e. 100% = complete gain learning; 0% = gain not leaned, error ratio is 3x larger than baseline error ratio). Learning percentages on RVF trials plateaued before the test phase in all groups (Fig. 2. B.), were moderately high in all groups (~80%), and did not significantly differ across groups (F (1,52) = 0.06, p = 0.807; Fig. 2. B. box plot). Thus, the greater transfer percentage found in the EPF groups could not be ascribed to better learning of the gain change.

### Right hand no visual feedback learning did not differ across groups

Learning percentages on RnoVF trials (Fig. 2. C.) were markedly worse than those seen for RVF, which was expected because visual feedback was not available to aid performance. Importantly, during the test phase RnoVF learning percentages did not differ across EPF and CVF groups (F(1,52 = 1.764, p = 0.19; Fig. 2. C. box plot). RnoVF performance at test gives an indication of how well the forward model has been updated during the learning phase. As such, if the EPF group had performed significantly better than the CVF group on these trials, one could conclude that that forward model learning was more pronounced with EPF. However, this was not the case and only the amount of intermanual transfer was found to be improved by EPF.

### No difference observed between Abrupt and Gradual perturbation schedules

No significant differences were observed when comparing abrupt and gradual perturbation schedules for all measures of RT and absolute error ratio. When comparing RVF RT at baseline and test, there was no significant main effect of perturbation schedule (F(1,52) = 0.001, p = 0.975) and no significant interaction between visual feedback type and perturbation schedule (F(1,52) = 1.242, p = 0.270). There was also no interaction between phase and perturbation schedule (F(1,52) = 1.742, p = 0.193), and no significant visual feedback type x perturbation schedule x phase interaction (F(1,52) = 0.013, p = 0.909). When baseline and test were considered separately, at baseline there was no significant main effect of perturbation schedule (F(1,52) = 0.181, p = 0.673) or perturbation x visual feedback type interaction (F(1,52) = 0.993, p = 0.324), and at test there was no significant main effect of perturbation schedule (F(1,52) = 0.152, p = 0.699) or perturbation x visual feedback type interaction (F(1,52) = 1.274, p = 0.264).

Likewise, transfer percentages at test did not significantly differ according to perturbation schedule (F(1,52) = 0.022, p = 0.883), and there was no significant perturbation schedule x visual feedback type interaction (F(1,52) = 0.003, p = 0.958). Baseline transfer percentage did not show a main effect of perturbation schedule (F(1,52 = 1.665, p = 0.203), nor a significant perturbation schedule x visual feedback type interaction (F(1,52 = 0.529, p = 0.471). RVF learning percentage also did not show a main effect of perturbation schedule (F (1,52) = 0.786, p = 0.380), nor a significant perturbation schedule x visual feedback type interaction (F (1,52) = 0.148, p = 0.720). Likewise, for RnoVF, learning percentages did not show a main effect of perturbation schedule (F(1,52 = 0.262, p = 0.611), nor a significant perturbation schedule x visual feedback type interaction(F(1,52 = 0.376, p = 0.542).

### Experiment 2 results

The purpose of Experiment 2 was to determine whether EPF was associated with smaller aftereffects than CVF, since smaller aftereffects have been associated with strategy use (Benson et al. 2011; Haith et al. 2015; Morehead et al. 2015). This was confirmed by assessing noVF trials after a sudden increase in visuomotor gain, which raised the gain back to baseline levels, following an extended period of adaptation to a gradually introduced lower level of visuomotor gain (Fig.3. A & B.). On noVF trials, the degree of overshoot was significantly larger for the CVF compared to the EPF group (Mn = 0.58, SD = 0.31 vs Mn = 0.26, SD = 0.21; t (27 = 3.247, p = 0.003, Cohen’s d = 1.2), indicating a larger aftereffect in the CVF group. Such aftereffects are hypothesised to be attenuated when participants use a strategy rather than adaptation to maintain performance during learning. On this account, participants in the EPF group displayed smaller aftereffects because they made greater use of a cognitive strategy throughout the task.

**Figure 3.**
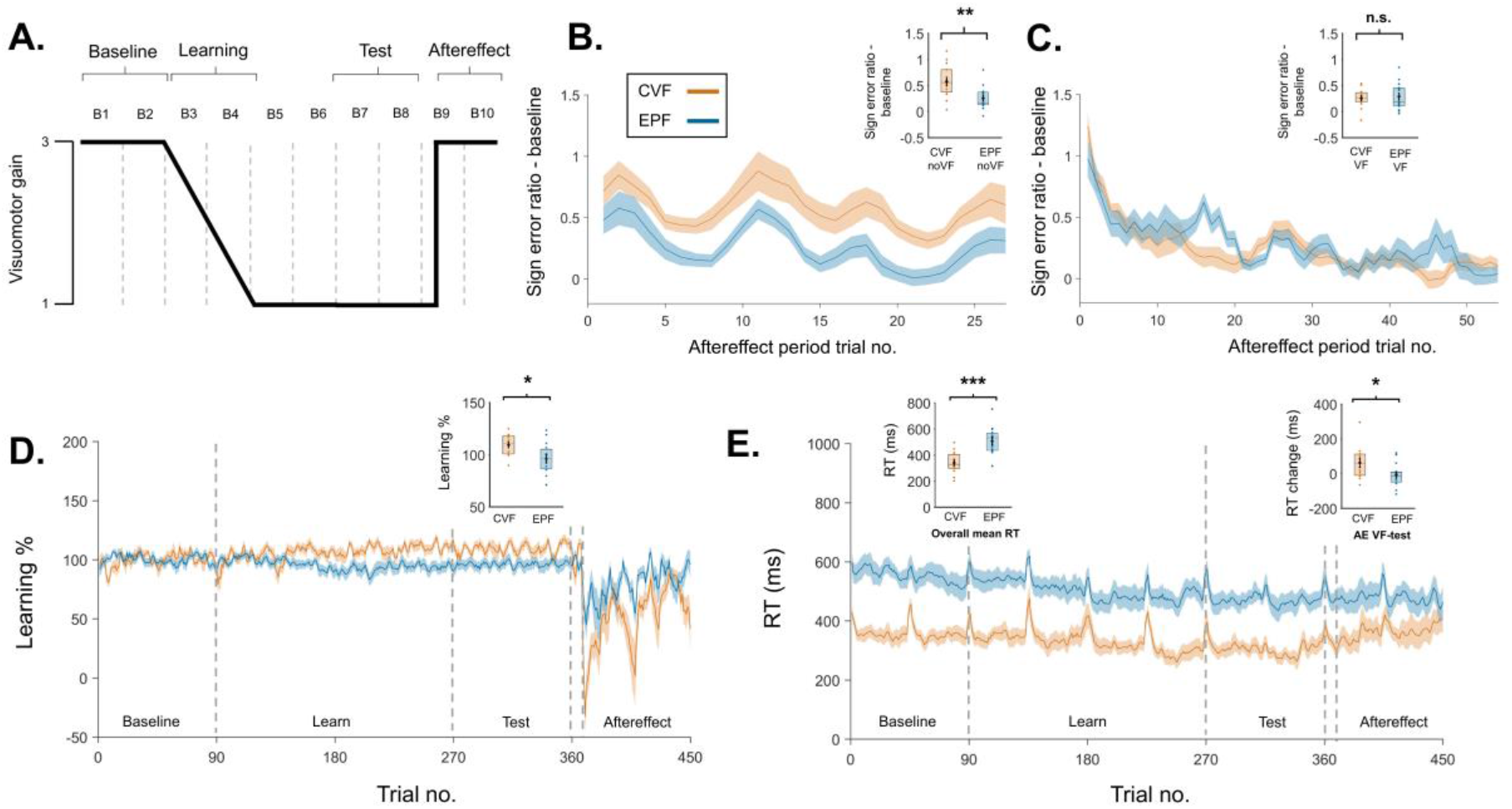
Design and results of Experiment 2. **A.** Design of Experiment 2 showing how visuomotor gain changed across blocks (B1-B10). On the 10^th^ trial of B9 there was sudden gain change which returned the gain to the baseline level. During this aftereffect phase trials alternated between 2 VF trials followed by 1 noVF trial. **B.** Signed error ratio after subtracting baseline values for noVF trials in the aftereffect phase, showing larger aftereffect for CVF than EPF groups. Box plot shows that the degree of overshoot was significantly higher for the CVF group relative to the EPF group, indicative of a greater aftereffect (** p <0.01). **C.** Signed error ratio after subtracting baseline values for VF trials in the aftereffect phase for CVF and EPF groups. Insert shows that there was no significant difference between the two groups (n.s. = not significant). **D.** RVF learning percentage across entire experiment for CVF and EPF groups. Note that both groups were able to maintain performance accuracy close to 100% from the baseline to the test phase. Insert shows that learning % at test was significantly higher in the CVF group compared to the EPF group (* p <0.05). **E.** Mean RT across entire experiment for CVF and EPF groups. Left box plot shows that overall RT was significantly slower in the EPF group compared to the CVF group. Right box plot shows that the CVF group increased their RT on VF trials from the test phase to the aftereffect phase to a greater extent than the EPF group (* p<0.05, ***p<0.001).

On VF trials participants in both groups were able to rapidly reduce their degree of target overshoot and there were no significant differences between the CVF and EPF groups (Mn = 0.26, SD = 0.18 vs Mn = 0.29, SD = 0.25; t (27) = −0.395, p = 0.696; Fig 3. C.).

We also examined how well the initial gradual gain decrease was learned. The learning percentage, based on the absolute error ratio, was close to 100% for both groups (Fig. 3. D.), but was significantly higher in the CVF group compared to the EP group (CVF Mn = 109.24%, SD = 10.09% vs EPF Mn = 95.97%, SD = 15.44%; t (27) = 2.719, p = 0.011, Cohen’s d = 1.02). Thus both groups learned the gradual gain change, but participants in the CVF group actually performed slightly better at test than baseline, while those in the EPF group performed slightly worse at test than baseline.

We replicated the finding from Experiment 1 that EPF was associated with longer RT than CVF (Mn = 506.14ms, SD = 113ms vs Mn = 340.25ms, SD = 81.3ms; t (27) = 4.509, p <0.001, Cohen’s d = 1.69; Fig. 3. E. left box plot), again consistent with the hypothesis that EPF promotes strategy use. As with Experiment 1, there was a general tendency for participants to respond faster from baseline to test, but this this speeding of responses did not differ between CVF and EPF groups (Mn = −53.08ms, SD = 53.98ms vs Mn = −83.67ms, SD = 41.73ms; t (27) = 1.714, p = 0.098).

Interestingly, participants in the CVF group tended to increase RT after the sudden gain change (aftereffect phase) on VF trials, while those in the EPF groups maintained their RT (Mn = 61.95ms, SD = 90.93ms vs Mn =−7.68ms, SD = 65.90ms; t (27) = 2.373, p = 0.025, Cohen’s d = 0.88; Fig. 3. E. right box plot). Indeed, by the end of the aftereffect phase group mean CVF RT rose to be similar to EPF RT. These results might indicate that the sudden gain change resulted in a greater reliance on strategy use in the CVF group. When questioned at the end of the experiment all participants in both groups reported being aware of the sudden gain increase (aftereffect phase), whilst remaining unaware of the earlier gradual gain decrease.

## Discussion

Encountering a change in the environment, humans can maintain their motor performance by either adapting their sensorimotor representation or by using a cognitive strategy to compensate for the change (Krakauer et al. 2019; Schween et al. 2020). We examined if elements of the design of a visuomotor task can facilitate a cognitive strategy, and whether this in turn enhances the intermanual transfer of learning. Across two experiments, when the visual feedback of our ballistic force production task was restricted to the endpoint, reaction times increased, suggesting a greater reliance on a cognitive strategy to solve the task (Haith et al. 2015; Klapp 1995). Following this pattern, intermanual transfer was facilitated in the endpoint feedback condition, indicating that a cognitive strategy can facilitate effector independent learning. We confirmed that EPF promoted strategy use via a second experiment which found reduced aftereffects relative to CVF.

Restricting visual feedback to the endpoint of the task (EPF) has been shown to facilitate cognitive control (Hinder et al. 2008; Taylor et al. 2014). During prism adaptation EPF may enhance the generalization of learning across effectors (Làdavas et al. 2011) and intermanual transfer (Cohen 1967), because it promotes greater strategic control than continuous visual feedback. Our results support the view that EPF encourages strategic learning because it involves a single performance error signal pertaining to goal completion. It therefore encourages learning at the planning stage, above the level of the control policy (McDougle et al. 2016). Strategic learning and motor adaptation have been suggested to involve dissociable brain networks (Jahani et al. 2020). The supplementary motor area is central to strategic control and bimanual tasks (Serrien et al. 2002), making it a likely candidate region for the processing of EPF and the associated generalizable learning we observed.

Cognitive strategies have always been considered more time consuming than motor adaptation (Fitts and Posner 1967). During an isometric visual rotation task, EPF was associated with slower RT, and the introduction of perturbations selectively slowed responses further under conditions of EPF (Hinder et al. 2008). Other studies have observed a relationship between cognitive strategy and longer RT (Benson et al. 2011; Fernandez-Ruiz et al. 2011; Saijo and Gomi 2010). Our results suggest that longer responding in our task can indicate strategic control throughout the task and that this form of learning can enhance performance when flexible responding is required.

In our task RT was not manipulated independently of visual feedback type, meaning that other interpretations of the prolonged RT in the EPF group, such as task difficulty, could not be completely excluded. It was therefore necessary to verify that EPF did indeed promote strategy use. In a second experiment we found that, having learned a gradual decrease of visuomotor gain, participants tended to overshoot the target after the gain suddenly increased back to baseline levels. These aftereffects in response to the “switching off” of a perturbation were consistent with those seen during reaching tasks (Galea et al. 2011; Kitago et al. 2013; Taylor et al. 2013; Taylor and Ivry 2011). Importantly, we found that aftereffect amplitude was markedly reduced in the EPF group relative to the CVF group, consistent with previous reports finding reduced aftereffects for EPF (Hinder et al. 2008; Barkley et al. 2014). Aftereffects are a hallmark of motor adaptation and reductions in aftereffects have previously been found to be caused by the use of cognitive strategies (Benson et al. 2011; Haith et al. 2015; Morehead et al. 2015). As such, enhanced intermanual transfer of learning associated with EPF can be ascribed to greater strategy use.

Changing the perturbation schedule from gradual to abrupt did not increase the RT, an indicator of strategy use, and consequently did not facilitate intermanual transfer of learning. Previous work indicated abrupt gain changes enhance intermanual transfer of learning via the facilitation of cognitive strategies (Malfait and Ostry 2004; Werner et al. 2019), but opposite results also exist (Taylor et al. 2011; Wang et al. 2011a, 2015). Since this strategy use is assumed to be due to the abrupt change causing the perturbation to reach explicit ‘awareness’ (Bouchard and Cressman 2021; Werner et al. 2019), difficulty in setting the size of the abrupt change and inter-individual variability in change sensitivity may explain the null-effect in the present study. In Experiment 1 abrupt gain changes were possibly also less salient than normal due to the constant switching between response hands. Indeed, in Experiment 2 we observed some evidence that that an abrupt change in gain (aftereffect phase) could produce some gradual slowing of RT in the CVF group, consistent with a slight increase in strategy use at the end of the task.

We found transfer percentages of ~85% for EPF, whereas previous motor tasks report ~25% (Balitsky Thompson and Henriques 2010; Sainburg and Wang 2002; Wang et al. 2011a; Wang and Sainburg 2004a). The disparity may be because of greater strategy use in our task, but it is also likely because transfer was assessed continuously, which has been shown to increase transfer rates from the 25% seen in blocked designs to above 50% (Taylor et al. 2011). Additionally, we used transfer from RnoVF to LnoVF trials, which controlled for task difficulty across conditions, but inherently gives larger transfer values than comparing to visual feedback trials. Caution is therefore required when comparing transfer rates across paradigms. We also only tested transfer from the right to the left hand. Several studies have reported that transfer is reduced from the non-dominant to the dominant arm (Criscimagna-Hemminger et al. 2003; Wang and Sainburg 2004b), while others have found no such asymmetry (Poh et al. 2016; Stockinger et al. 2015; Wang et al. 2011b). Future work is needed to address how left to right transfer works during force production tasks.

Motor learning occurs at multiple levels of the control hierarchy (Hikosaka et al. 1999), with movement planning involving effector dependent and independent brain regions (Gallivan et al. 2011). Intermanual transfer of learning has generally been assumed to be achieved by updating such effector independent motor representations (Ruddy and Carson 2013). However, an effector independent cognitive control strategy, such as re-aiming (Benson et al. 2011), can achieve the same result. Future models of intermanual transfer need to consider the role of cognitive strategies.

In conclusion, our results add to the growing body of literature showing that elements of the task environment, such as the type of visual feedback available, can alter the balance between cognitive strategies and motor adaptation. The involvement of a cognitive strategy enhances intermanual transfer of learning. This greater generalization may result from strategic learning being related to movement planning, and as such being located above the control policy in the motor hierarchy.

## Conflict of interest statement

The authors declare no conflicts of interest

## Author contributions

JDH, PH, HG, SB, YI and NH conceived of the presented idea. JDH and NH designed the experiment. JDH and NH collected the data. Data was analysed by JDH and NH with support from HG. JDH and NH wrote the manuscript. All of the authors read and commented on the manuscript.

## Acknowledgements

None

## Funding

JDH was supported by a UCL-NTT Impact studentship and by a NICT Internship Trainee Program. NH is supported by Japan Society for the Promotion of Science (Kakenhi 26119535, 18H01106). YI and NH are supported by ERATO (JPMJER1801).

## Notes

### Competing Interest Statement

The authors have declared no competing interest.

